# Methods for Automatic Reference Trees and Multilevel Phylogenetic Placement

**DOI:** 10.1101/299792

**Authors:** Lucas Czech, Alexandros Stamatakis

## Abstract

**Motivation:** In most metagenomic sequencing studies, the initial analysis step consists in assessing the evolutionary provenance of the sequences. Phylogenetic (or Evolutionary) Placement methods can be employed to determine the evolutionary position of sequences with respect to a given reference phylogeny. These placement methods do however face certain limitations: The manual selection of reference sequences is labor-intensive; the computational effort to infer reference phylogenies is substantially larger than for methods that rely on sequence similarity; the number of taxa in the reference phylogeny should be small enough to allow for visually inspecting the results.

**Results:** We present algorithms to overcome the above limitations. First, we introduce a method to automatically construct representative sequences from databases to infer reference phylogenies. Second, we present an approach for conducting large-scale phylogenetic placements on nested phylogenies. Third, we describe a preprocessing pipeline that allows for handling huge sequence data sets. Our experiments on empirical data show that our methods substantially accelerate the workflow and yield highly accurate placement results.

**Implementation:** Freely available under GPLv3 at http://github.com/lczech/gappa.

**Supplementary Information:** Supplementary data are available at *Bioinformatics* online.

## 1 Introduction

High-throughput DNA sequencing technologies have revolutionized biology by transforming it into a data-driven computational discipline (Escobar-Zepeda *et al.*, 2015). Next Generation Sequencing (NGS) methods now allow for studying microbial samples directly extracted from their environment (Edwards and Holt, 2013). For each sample, these methods yield a set of short, anonymous DNA sequences, so-called reads. A typical task in such studies is to identify and classify the reads by relating them to known reference sequences, either taxonomically or phylogenetically.

Conventional methods based on sequence similarity are fast and work reasonably well if the reads are similar enough to the reference sequences, that is, if they represent species that are closely related to known species. However, they might *not* yield the most closely related species (Koski and Golding, 2001). This is particularly true for environments where available reference databases do not exhibit sufficient taxon coverage (Mahé *et al.*, 2017). As insufficient taxon coverage cannot be detected by methods that are based on sequence similarity, they can potentially bias downstream analyses.

So-called phylogenetic (or evolutionary) placement methods (Matsen *et al.*, 2010; Berger *et al.*, 2011; Barbera *et al.*, 2018) provide a more accurate means for identifying reads. Instead of relying on sequence similarity, they identify reads based on a phylogenetic tree of reference sequences. Thereby, they can incorporate information about the evolutionary history of the species under study.

In short, phylogenetic placement calculates the most probable insertion branches for a query sequence (QS) on a given reference tree (RT). For metagenomic studies, the QSs are the reads from the environmental samples. First, the QSs are aligned against the reference alignment of the RT. For a given QS and a given branch in the RT, the QS is inserted as a new tip into the branch; the affected branch lengths are then re-optimized; the likelihood score of the tree is evaluated; and the QS is removed again from the branch. This process yields a so-called *placement* of the QS for every branch of the RT, that is, an optimized position on the branch, along with a likelihood score for the entire RT. These likelihood scores are then transformed into probabilities to quantify the uncertainty of the QS placement into the respective branch. This placement process is repeated independently for each QS on the original RT. Phylogenetic placement thus yields a mapping of each QS to all branches of the RT, along with a probability for each placement of a QS on a specific branch.

Phylogenetic placement is particularly helpful for studying new, unexplored environments, for which no closely related sequences exist in reference databases (e.g., (Mahé *et al.*, 2017)). However, the selection of suitable reference sequences for inferring the RT constitutes a challenge for studying such environments. Furthermore, conducting phylogenetic placements requires a higher computational effort with respect to the placement algorithms *per se*, but also the pre- and post-processing, than, for instance, similarity based methods. Nonetheless, existing placement algorithms are being increasingly used and cited. Due to the continuous advances in molecular sequencing, existing placement methods as well as respective pre- and post-processing tools have already reached their scalability limits.

## 2 Methods

Here, we introduce methods to overcome the aforementioned limitations, that is, to (1) automatically obtain a high quality reference tree for phylogenetic placement, (2) split up the placement process into two steps using smaller phylogenies, and (3) accelerate the computation of placements via appropriate data pre-processing approaches.

### 2.1 Automatic Reference Trees

#### 2.1.1 Motivation

Molecular environmental sequencing studies, particularly those that aim to conduct phylogenetic placement, often rely on a set of manually selected and aligned reference sequences to infer an RT (Tedersoo *et al.*, 2014; de Vargas *et al.*, 2015; Mahé *et al.*, 2017; Thompson *et al.*, 2017). Creating and maintaining databases of such reference sequences constitutes a labor-intensive and potentially error-prone process. Moreover, this approach is impractical for samples from large domains or samples obtained from unexplored environments. Lastly, even if a large RT is available, the visualization of placements on such an RT might be confusing and thus hard to interpret.

The reference tree (RT) used for phylogenetic placement should ideally (a) cover all major taxonomic groups that occur in the QSs, (b) use high-quality error-free reference sequences, and (c) not be too large to allow for unambiguous visualization and interpretation. These criteria can be met for small datasets by manually selecting curated sequences from databases. For large and taxonomically diverse samples one key challenge is that sequence databases such as Greengenes (DeSantis *et al.*, 2006), Unite (Abarenkov *et al.*, 2010), PR2 (Guillou *et al.*, 2012), Ez-Taxon (Kim *et al.*, 2012), Silva (Quast *et al.*, 2013), and RDP (Cole *et al.*, 2014) maintain reference collections of thousands to millions of taxonomically annotated sequences. Therefore, one needs to appropriately sub-sample sequences such that the RT can be inferred in reasonable time *and* and sufficiently covers the diversity of the sample.

To this end, we present a computationally efficient approach for constructing sequences from large databases to infer an RT. This RT is then used for conducting phylogenetic placement analyses. Our approach identifies sets of sequences that are similar to each other based on their entropy. It subsequently reduces the sequences in these sets to a predefined number of consensus sequences that can in turn be used to infer the RT. The input of our method is a database of aligned sequences of known species including their taxonomic labels.

#### 2.1.2 Sequence Entropy

First, we define a measure to quantify the ensemble similarity of a set *s* of sequences, based on their entropy (Shannon and Weaver, 1951). Variants of sequence entropy have been used before in numerous biological and phylogenetic contexts, for example, to asses the information content of sequences (Schmitt and Herzel, 1997; Vinga, 2014), or to measure substitution saturation (Xia *et al.*, 2003). Here, we use entropy for alignment sites, that is, we define the entropy (uncertainty) *H* at alignment site *i* as

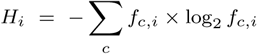

where *c* ∈ {A, C, G, T, −} is the set of nucleotide states including gaps, and *f*_*c,i*_ is the frequency of character *c* at site *i* of the alignment. Including gaps (-) in the summation reduces the contribution of sites that contain a large fraction of gaps. Their contribution is weighed down as all standard phylogenetic inference tools model gaps as undetermined states, that is, they do not contribute anything to the likelihood score. The entropy is 0 for sites that only contain a single character. It increases the more different characters an alignment site contains, *and* the more similar their frequencies are. Its maximum occurs if all characters appear with the same frequency (each of them 20%). Note that we also treat ambiguous characters as gaps. As only 0.008% of the non-gap characters in our test database (Silva) are ambiguous, their influence is negligible. Ambiguous characters could however be incorporated by using fractional character counts.

Finally, the total entropy of a set *s* of aligned sequences is simply the sum over all per-site entropies: *H*(*s*) = *Σ_i_. H_i_*. We use this entropy to quantify the ensemble similarity of a set of sequences. This can be regarded as an information content estimate of the sequences.

#### 2.1.3 Sequence Grouping

The goal of this step is to group the sequences of a database into a given target number of groups/sets, such that the groups reflect the diversity of the sequences in the database. We use the taxonomy to identify potential candidate groups of sequences that could be represented by a consensus sequence. We interpret a taxonomy as a sequence labeling, where similar sequences have related labels. Thus, a taxonomy represents a pre-classification of similar sequences that can be exploited to group them.

For a clade *t* of the taxonomic tree, we denote by *H*(*t*) the entropy of all sequences that belong to that clade, including all sequences in its sub-clades, that is, its lower taxonomic ranks. Clades with low entropy imply that they contain highly similar sequences that can in turn be represented by a consensus sequence without sacrificing too much diversity. Inversely, clades with high entropy contain diverse sequences, implying that a consensus sequence is not likely to sufficiently capture the inherent sequence diversity. It is thus better to expand these clades and construct separate consensus sequences for their respective sub-clades. An example is shown in Figure 1. As the clade structure of a taxonomy forms a tree, this criterion can then be applied recursively, as shown in Algorithm 1.

**Figure 1:**
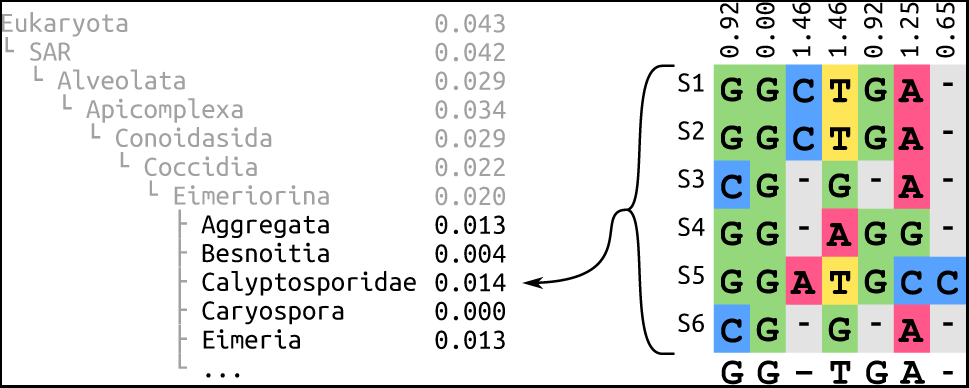
Entropy and consensus sequence of a taxonomic clade. The left hand side shows the exemplary clade *Eimeriorina* in its taxonomic context, listing its super- and sub-clades with the normalized entropy of their respective sequences. The right hand side is an excerpt from the alignment of six sequences that belong to the *Calyptosporidae* sub-clade. At its top, the per-site entropies for the alignment columns are shown. At the bottom, the majority rule consensus sequence is shown, which is used to represent the sub-clade.

The algorithm works as follows: We initialize a list of candidate clades with the highest ranking clades that we want to consider. In the most general case, these can be “Archaea”, “Bacteria”, and “Eukaryota”. We then select the most diverse candidate clade, that is, the clade *t* whose sequences exhibit the highest entropy *H*(*t*). This clade is then expanded, and we do not consider it as a potential candidate for building a consensus sequence. The high entropy clade is then removed from our list and its immediate sub-clades are added as new candidates to the list. Finally, the current count of how many candidates we have already selected is updated accordingly. By expanding clades with high entropy, we descend into the lower ranks of the taxonomy. On average, this decreases the entropy, because low ranking clades generally tend to contain more similar sequences. This process is repeated until our list contains approximately as many candidate clades as the desired target count of reference sequences, which is provided as input. As the sizes of expanded clades can vary substantially, the target count cannot always be met exactly. In our tests, the average deviation was 0.2%, as shown in Supplementary Table S1.

##### Algorithm 1 Taxonomy Expansion

~~~
*Candidates ←* list of highest ranking clades
*TaxaCount ←* size of *Candidates*
**while** *TaxaCount < TargetCount* **do**
        *MostDiverse ←* arg max _*t∈Candidates*_ *H*(*t*)
        remove *MostDiverse* from *Candidates*
        add sub-clades of *MostDiverse* to *Candidates*
        *TaxaCount ← TaxaCount −*1 + size of *MostDiverse*
**return** *Candidates*
~~~

Given this list of clades from different taxonomic ranks, we can now compute the consensus sequences. For each clade, all sequences in that clade and its sub-clades are used to construct a consensus sequence, which represents the clade diversity, and serves as the reference sequence for that clade. A simple per-site majority rule consensus (May, 1952) works well, but we also assessed alternative methods; see Supplementary Figure S2 for details. The above process yields a set of consensus reference sequences which capture the diversity of distinct taxonomic clades.

#### 2.1.4 Inferring a Reference Tree

Once we have identified the consensus sequences, which are already aligned to each other, we can use them to infer a maximum likelihood tree, which we call an *Automatic Reference Tree* (ART). As each consensus sequence is associated with a taxonomic clade, the corresponding taxonomic path can be used to label the tips of the tree. Note that since clades with low entropy might not be expanded, the tip labels do not necessarily correspond to species or genera. Also, the ART will not necessarily be congruent to the taxonomy.

An ART obtained via our method satisfies all criteria we listed: (a) All taxonomic groups occurring in the QSs can be covered by using a suitable taxonomy as input. (b) By using consensus sequences, potential sequencing errors can be alleviated. (c) The size of the ART can be specified by the user. However, using consensus sequences may obscure the degree of sequence diversity in sub-clades, which in turn can affect the accuracy of subsequent phylogenetic placements on that tree. Our algorithm as described here can not fully compensate for this. We present a method to address this issue in the next Section.

### 2.2 Multilevel Placement

When conducting phylogenetic placement, the computationally limiting factors are (i) the number of QSs to be placed (addressed in the next section) and (ii) the size of the RT (number of taxa) and corresponding alignment length (addressed below). Using RTs with more taxa increases the phylogenetic resolution of the placements, at the cost of increased computational effort for inferring the RT, aligning the QSs, and placing the QSs. Furthermore, longer reference alignments (if appropriate data is available) are required to accurately infer large trees under the maximum likelihood criterion (Yang, 1994), thus further increasing the computational costs. Lastly, placement on large trees that comprise reference sequences with high evolutionary distances can reduce placement accuracy (Mirarab *et al.*, 2012). Thus, using a large number of reference sequences is not always desirable in practice.

To address this issue, we present an approach called *Multilevel* or *Russian Doll* Placement, which is summarized in Figure 2. Instead of working with one large RT comprising *all* taxa of interest, we use a smaller, but taxonomically broad backbone tree (BT) for pre-classifying the QSs (first level), and a set of refined clade trees (CTs) for the final, more accurate placements (second level). These CTs comprise the reference sequences that are of interest for a particular study. Each CT is associated with the set of branches of a specific BT clade.

**Figure 2:**
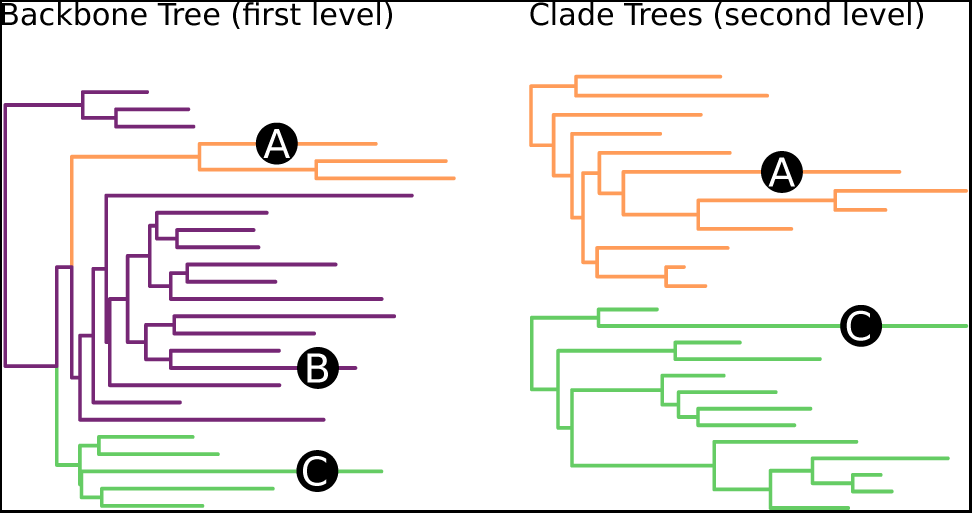
Multilevel Placement. The left shows a backbone tree (BT); the right shows two clade trees (CTs) in orange and green. Branches in the BT that are associated with a CT are marked in its color. The trees “overlap” each other, meaning that each CT is represented by multiple branches in the BT. Three sequences A, B and C are placed on the BT, which is the first level. A and C are placed on branches associated with a CT. Hence, their second level placement is conducted on the respective CT. B is placed on a branch that is not associated with any CT, and thus not used in the second level.

The method then works in three steps:

1. Align and place the QSs using the BT (first level).
2. For each CT, collect the QSs that are placed on the BT branches associated with the CT.
3. Align and place these QSs again, using their specific CTs (second level).

While this approach requires some additional bookkeeping, the total computational cost is reduced, because the QSs do not have to be placed on all branches of all CTs. Furthermore, this method allows for fine-grained control over the clades of interest at both placement levels:

Firstly, the BT provides a means for phylogenetically informed sequence filtering – that is, to identify and remove “spurious” QSs. Sequences with low similarity to known references are often removed in environmental sequencing studies. However, using sequence similarity as a filter criterion can remove too many QSs, particularly when studying new, unexplored environments (Mahé *et al.*, 2017). By using phylogenetic placement as a filter instead, substantially more sequences can be retained for downstream analyses. Only the QSs that are placed onto the inner branches of the BT, that is, branches with no associated CT, are omitted at the second placement level.

Secondly, using specific clade trees for lower level taxonomic clades offers the phylogenetic resolution that is necessary for downstream analyses and for biological reasoning. It is, for example, possible to use manually curated “expert” trees for each clade of interest.

In this setup, the BT is only used for pre-classification, and can, for example, use our Automatic Reference Tree method. The aforementioned issue of obscured diversity in sub-clades can be circumvented by “overlapping” the CTs with the BT. That is, a CT can be associated with several branches of the BT, so that placements on each of these BT branches are collected and placed onto the same CT. See Figure 2 and Supplementary Figure S3 for examples. We recommend to ensure that the branches of the BT that are associated with one CT are monophyletic, meaning that there is one split that separates these branches from the rest of the BT. This can be achieved by inferring the BT with a high-level constraint that maintains the monophyly of the CTs. It ensures phylogenetic consistency between the BT and the CTs, and improves the accuracy of the first placement level, as shown in Section 3.3. Lastly, it is also possible to use more than two levels, which might become necessary when working with RTs and datasets even larger than what is currently available.

### 2.3 Data Preprocessing for Phylogenetic Placement

Apart from the RT size, handling the sheer number of QSs also induces computational limitations for conducting phylogenetic placements. Most metagenomic studies publish their data in unprocessed formats. Those data often contain duplicates of exactly identical sequences, both *within* and *across* samples. Identical sequences are however treated the same in phylogenetic placement algorithms and therefore induce unnecessary computational overhead. Furthermore, sample sizes, that is, the number of sequences per sample, can vary by several orders of magnitude. If the placement algorithm is parallelized over samples, this leads to an uneven load balance across compute nodes.

In order to solve these issues, that is, reduce computational cost and achieve good load balancing, one can pre-process the sequences with our gappa tool. First, sequences are de-duplicated across all samples and fused into chunks of equal size. The chunk size should be chosen to allow aligning and placing a chunk within wall time on the intended hardware; we recommend chunk sizes of 50 000 or larger. Our tool assigns an identifier to each unique sequence, and computes a list of abundance counts for each sequence in a sample. Given an RT and its underlying alignment, the QS chunks are then aligned to the reference multiple sequence alignment, using programs such as PaPaRa (Berger and Stamatakis, 2012) or hmmalign (Eddy, 1998), and subsequently placed on the RT, for example by pplacer, RAxML-EPA or EPA-ng (Matsen *et al.*, 2010; Berger *et al.*, 2011; Barbera *et al.*, 2018). The resulting per-chunk placement result files in combination with the per-sample abundance counts can then be parsed and analyzed by gappa to generate final per-sample placement files, containing a placement for each sequence in the original sample.

This approach allows to analyze datasets that are orders of magnitude larger than in previous published studies. For example, in 2012, an analysis of Bacterial Vaginosis (BV) data placed 15 060 unique sequences on an RT with 796 tips (Srinivasan *et al.*, 2012). Using a prototype of gappa, we were able to analyze 10 567 804 unique sequences from neotropical soils with an RT comprising 512 taxa (Mahé *et al.*, 2017). To demonstrate the scalability of our methods for this paper, we analyzed datasets with up to 63 221 538 unique sequences from the Human Microbiome Project (HMP) (Huttenhower *et al.*, 2012; Methé *et al.*, 2012), using RTs with up to 2059 tips. This corresponds to a computational effort that is four orders of magnitude greater than for the BV study.

## 3 Results

To test the automatic reference tree (ART) method, we used the “SSU Ref NR 99” sequences of the Silva database (Quast *et al.*, 2013) version 123.1 and the corresponding taxonomic framework (Yilmaz *et al.*, 2014). The database contains 598 470 aligned sequences from all three domains of life, classified into 11 860 distinct taxonomic labels.

We constructed four sets of consensus sequences from the Silva database: a *General* set (“all of life”), as well as separate sets for the domains *Archaea*, *Bacteria*, and *Eukaryota*. For each set except the *Archaea*, the recursive expansion of taxonomic clades was applied to obtain approximately 2000 (*General*) and 1800 (*Bacteria*, *Eukaryota*) consensus reference sequences. This is large enough to cover the diversity well, while still being computationally feasible for the subsequent steps. The *Archaea* taxonomy in Silva is smaller, containing 248 taxa at *Genus* level, which is the lowest level in their taxonomy. Hence, the *Archaea* tree also comprises 248 taxa. Furthermore, in the three domain-specific trees, we included sequences at the *Phylum* level of the respective two other domains, to ensure that our methods also work with outgroups. An overview of the tree sizes is shown in Supplementary Table S1. We then inferred constrained and unconstrained maximum likelihood trees for the consensus sequences. The constrained trees comply with the Silva taxonomy, and are used to assess how taxonomic constraints affect the phylogenetic placement and the subsequent analyses. Details are provided in Supplementary Section 1, which also discusses differences between the constrained and unconstrained trees. Details of the trees are shown in Supplementary Table S3; Supplementary Figure S3 shows the unconstrained *Bacteria* tree as an example.

In total, our setup yields eight distinct RTs for evaluation: the *General* tree, the three domain trees, and the respective taxonomically constrained variants.

### 3.1 Accuracy

Here, we assess how using our ART affects phylogenetic placement accuracy. Each terminal branch of our RTs represents a consensus sequence, which is computed from species level sequences that share the same taxonomic label. We evaluate an RT by placing these species sequences onto the RT: Each species sequence is expected to be placed onto the branch leading to the consensus sequence that represents this particular species sequence. For example, sequences S1-6 in Figure 1 are represented by the consensus sequence for the *Calyptosporidae* clade. They are thus expected to be placed onto the *Calyptosporidae* branch in the RT.

We placed the respective subset of the Silva database species sequences onto each of the eight RTs. We quantify placement accuracy for a sequence by the distance to its expected placement branch. More precisely, we measured (a) the (discrete) number of branches between the actual placement and the expected branch, and (b) the (continuous) distance in branch lengths units. As a sequence can have multiple placement locations, the distances, are, in fact, weighted averages incorporating the placement probabilities (likelihood weights). The results for the four unconstrained trees are shown in Figure 3; Supplementary Figure S1 depicts the results for the constrained trees. Further details are provided in Supplementary Table S3.

**Figure 3:**
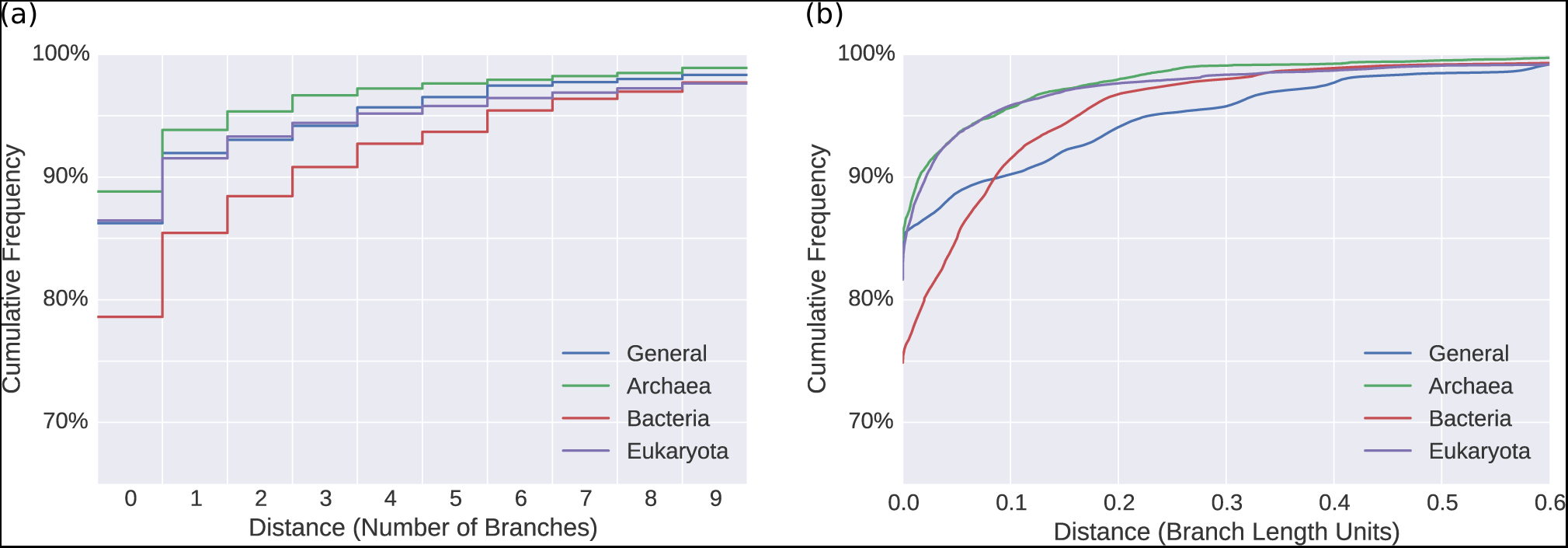
Weighted distances to expected edges for unconstrained trees. We evaluated the accuracy of our ARTs by placing sequences and measuring the weighted distances to their respective expected placement branches. The Figure shows the cumulative frequencies of number of sequences versus distances, measured (a) in number of branches and (b) in branch length units. In other words, it shows how many sequences are placed within a certain radius from their expected branches. For example, in (a), more than 85% of the sequences of the *Bacteria* (red) are placed within a radius of at most one branch from their expected branch, and in (b), more than 95% of the Eukaryota (purple) are within a radius of 0.1 branch length units from their expected branches.

Considering the size of the trees, most sequences are placed in close vicinity to their expected branches. This is corroborated by the short average distances reported in Supplementary Table S3. Furthermore, the average expected distance between placement locations (EDPL, Matsen *et al.*, 2010) is low, indicating that the placements of a specific sequence mostly cluster in a small neighborhood of the tree. We observed that errors occur mostly in parts of the tree with short branches, which might be explained by the inability of 16S SSU sequences to properly resolve certain clades (Janda and Abbott, 2007). Also, the placement likelihood differences are small between neighboring, short branches, such that the placement signal is fuzzy.

With 77% of the sequences placed exactly on their expected branch, the accuracy is generally lowest for the *Bacteria* tree. This might be because the *Bacteria* have the most sequences in Silva, and exhibit a high diversity. In the other three trees, more than 90% of the sequences are placed at most one branch away from their respective expected branch. The constrained trees (Supplementary Figure S1) exhibit similar placement accuracy. Particularly when using Multilevel Placement with overlapping RTs, placement differences of a few branches on the first level tree are acceptable, as they do not change the second level tree on which the sequence is placed. See Section 3.3 for details.

As outlined in the method description, we represent clade diversity via majority rule consensus sequences. To assess the impact of the consensus method, we repeated the above evaluation, using two alternative consensus methods, but found little difference between the methods (see Supplementary Figure S2).

### 3.2 Empirical Datasets

ARTs are intended for obtaining phylogenetic placements of environmental sequences. As the true evolutionary history of such sequences is unknown, we can not repeat the previous accuracy tests on empirical environmental datasets. Instead, we assess if the ARTs yield meaningful quantitative results for typical post-analysis methods. To this end, we placed two empirical metagenomic read datasets on our unconstrained *Bacteria* tree. To asses the placement results obtained from the ART, we performed Squash Clustering and Edge PCA (Matsen and Evans, 2011) post-analyses on the placement results (see Supplementary Section 2 and Supplementary Figures S4 and S5 for details). The results reveal that the ART reproduces results of previous studies based on custom RTs. Furthermore, the ART is able to classify samples (e.g., healthy vs sick patients), at least to the extend that is expected from its phylogenetic resolution. That is, samples that only differ in placements at the species level cannot be classified using a broad, high-level tree such as our *Bacteria* tree. In order to obtain finer taxonomic resolution, it is thus necessary to either use an ART that contains more taxa, or to use our multilevel approach instead (see next Section).

### 3.3 Sub-clades and Multilevel Placement

We selected five bacterial clades to evaluate ART accuracy on smaller clades, as well as to assess some properties of the Multilevel Placement approach. Supplementary Figure S3 shows the *Bacteria* tree with the five test clades highlighted.

First, using the sequences and taxonomies of these five clades, we built unconstrained and constrained ARTs. We then conducted the same accuracy analysis as explained before on these ten trees. That is, we placed the Silva sequences of the five clades onto their respective ART and evaluated distances to expected branches. Thereby, we evaluated the accuracy of these ARTs when used as second level clade trees. The results are shown in Supplementary Figure S6. The placement accuracy is slightly worse for the clade trees than for the eight comprehesive ARTs evaluated before. This is again likely due to 16S SSU sequences being unable to properly resolve lower taxonomic levels (Janda and Abbott, 2007).

Next, using the five clades, we evaluated the accuracy of the first placement level when conducting Multilevel Placement. So far, our evaluation focused on the distance from a sequence placement to its expected placement branch. For the first placement level on a backbone tree (BT), it is however more important that a sequence is placed into the correct clade. Thus, we used the unconstrained *Bacteria* BT again, and assessed how many sequences were placed in the clades shown in Figure S3. Of the 450 313 sequences in Silva in these clades, 98.0% were placed (most likely placement) into a branch of their corresponding clade. Thus, for multilevel placement, they will be assigned to the correct second level clade tree (CT). More specifically, the *Firmicutes* perform worst, as only 94.7% of the *Firmicute* sequences are placed into the corresponding clade. This can be explained by the high amount of paraphyletic branches of this clade, cf. Figure S6, which is a known issue (Parks *et al.*, 2018).

The sequences of the other four clades we tested achieve a clade identification accuracy exceeding 99%.

As mentioned before, a high level taxonomic constraint can improve the accuracy of placing a sequence into the correct BT clade. To show this, we inferred the *Bacteria* RT again, but used a *Phylum* level constraint that separates the five clades from each other and from the rest of the tree. All branches within the clades were resolved using maximum likelihood. The tree (not shown) is similar to the tree in Figure S3, but all five clades are now monophyletic. Using this tree, 99.3% of the sequences were placed into the correct clade. Particularly the accuracy for *Firmicutes* improved, yielding an accuracy of 99.5%.

Overall, our experiments show that the first level placement is highly accurate, even if an extremely diverse “all bacteria” backbone tree is used. The accuracy on the second level is slightly worse when using ARTs as CTs.

## 4 Discussion and Conclusion

We presented algorithms and software tools to facilitate and accelerate phylogenetic placement of large environmental sequencing studies.

The automatic reference tree (ART) method provides a means for automatically obtaining suitable reference trees by using the taxonomy of large sequence databases. Using the Silva database as a test case, we showed that it can be applied for accurately (pre-)placing environmental sequences into taxonomic clades. The method can also be used for rapid data exploration in environmental sequencing studies: An ART might be useful to obtain an overview of the taxa that are necessary to capture the diversity of a sequence dataset, without the substantial human effort and potential bias of manually selecting reference sequences. To capture clade diversity with finer resolution, for example for a second placement level, clade-specific ARTs can be inferred. If species-level resolution is required, we recommend that the sequences are inspected by an expert, as our automated approach inevitably suffers from errors in the database it is based on. We therefore recommend using Sativa (Kozlov *et al.*, 2016) to identify potentially mislabeled sequences in the database. One should keep in mind that phylogenetic placement does not necessarily provide resolution at the species level (Dunthorn *et al.*, 2014).

As we show, our multilevel placement method as well as the preprocessing pipeline accelerate the placement process without sacrificing accuracy. By first placing the query sequences on a broad backbone tree (BT), novel environments with sequences of unknown evolutionary origin can be classified without having to process a large tree comprising all taxa of interest. A second placement on a set of clade trees (CTs) provides sufficient resolution for biological interpretation. Placement accuracy can be further improved by inferring the BT with a high-level constraint that separates the clades of the CTs from each other and thus ensures monophyly of these clades.

All methods are implemented in our gappa tool, which is freely available under GPLv3 at http://github.com/lczech/gappa (see Supplementary Section 3 for an overview of the corresponding commands). All scripts and data used for this paper are available at http://github.com/lczech/placement-methods-paper.

## Acknowledgements

We thank **S. Srinivasan** and **E. Matsen** for providing the Bacterial Vaginosis dataset (Srinivasan *et al.*, 2012); **M. Dunthorn**, **G. Lentendu**, **P. Barbera** and **A. Kozlov** for their feedback on our methods and this manuscript.

## References

Abarenkov, K. et al. (2010). The UNITE database for molecular identification of fungi–recent updates and future perspectives.New Phytologist, 186(2), 281–285.

Barbera, P. et al. (2018). EPA-ng: Massively Parallel Evolutionary Placement of Genetic Sequences. bioRxiv.

Berger, S. and Stamatakis, A. (2012). PaPaRa 2.0: A Vectorized Algorithm for Probabilistic Phylogeny-Aware Alignment Extension. Technical report, Institute for Theoretical Studies, Heidelberg.

Berger, S. et al. (2011). Performance, accuracy, and web server for evolutionary placement of short sequence reads under maximum likelihood. Systematic Biology, 60(3), 291–302.

Cole, J. R. et al. (2014). Ribosomal database project: data and tools for high throughput rRNA analysis. Nucleic Acids Res, 42.

de Vargas, C. et al. (2015). Eukaryotic plankton diversity in the sunlit ocean. Science, 348(6237), 1261605–1261605.

DeSantis, T. Z. et al. (2006). Greengenes, a chimera-checked 16S rRNA gene database and workbench compatible with ARB. Applied and environmental microbiology, 72(7), 5069–5072.

Dunthorn, M. et al. (2014). Placing environmental next-generation sequencing amplicons from microbial eukaryotes into a phylogenetic context. Molecular Biology and Evolution, 31(4), 993–1009.

Eddy, S. R. (1998). Profile hidden Markov models. Bioinformatics, 14(9), 755—-763.

Edwards, D. J. and Holt, K. E. (2013). Beginner’s guide to comparative bacterial genome analysis using next-generation sequence data. Microbial informatics and experimentation, 3(1), 2.

Escobar-Zepeda, A. et al. (2015). The road to metagenomics: From microbiology to DNA sequencing technologies and bioinformatics. Frontiers in Genetics, 6(DEC), 1–15.

Guillou, L. et al. (2012). The Protist Ribosomal Reference database (PR2): a catalog of unicellular eukaryote small subunit rRNA sequences with curated taxonomy. Nucleic acids research, 41(D1), D597–D604.

Huttenhower, C. et al. (2012). Structure, function and diversity of the healthy human microbiome. Nature, 486(7402), 207–214.

Janda, J. M. and Abbott, S. L. (2007). 16S rRNA Gene Sequencing for Bacterial Identification in the Diagnostic Laboratory: Pluses, Perils, and Pitfalls. Journal of Clinical Microbiology, 45(9), 2761–2764.

Kim, O.-S. et al. (2012). Introducing EzTaxon-e: a prokaryotic 16S rRNA gene sequence database with phylotypes that represent uncultured species. International journal of systematic and evolutionary microbiology, 62(3), 716–721.

Koski, L. B. and Golding, G. B. (2001). The closest BLAST hit is often not the nearest neighbor. Journal of molecular evolution, 52(6), 540–2.

Kozlov, A. M. et al. (2016). Phylogeny-aware identification and correction of taxonomically mislabeled sequences. Nucleic Acids Research, 44(11), 5022–5033.

Mahé, F. et al. (2017). Parasites dominate hyperdiverse soil protist communities in Neotropical rainforests. Nature Ecology & Evolution, 1.

Matsen, F. A. and Evans, S. N. (2011). Edge principal components and squash clustering: using the special structure of phylogenetic placement data for sample comparison. PLOS ONE, 8(3), 1–17.

Matsen, F. A. et al. (2010). pplacer: linear time maximum-likelihood and Bayesian phylogenetic placement of sequences onto a fixed reference tree. BMC Bioinformatics, 11(1), 538.

May, K. O. (1952). A set of independent necessary and sufficient conditions for simple majority decision. Econometrica: Journal of the Econometric Society, pages 680–684.

Methé, B. A. et al. (2012). A framework for human microbiome research. Nature, 486(7402), 215–221.

Mirarab, S. et al. (2012). SEPP: SATé-Enabled Phylogenetic Placement. Biocomputing, pages 247–258.

Parks, D. H. et al. (2018). A proposal for a standardized bacterial taxonomy based on genome phylogeny. bioRxiv.

Quast, C. et al. (2013). The SILVA ribosomal RNA gene database project: improved data processing and web-based tools. Nucleic Acids Research, 41(D1), D590–D596.

Schmitt, a. O. and Herzel, H. (1997). Estimating the entropy of DNA sequences. Journal of theoretical biology, 188(3), 369–377.

Shannon, C. E. and Weaver, W. (1951). The Mathematical Theory of Communication. University of Illinois Press.

Srinivasan, S. et al. (2012). Bacterial communities in women with bacterial vaginosis: High resolution phylogenetic analyses reveal relationships of microbiota to clinical criteria. PLOS ONE, 7(6), e37818.

Tedersoo, L. et al. (2014). Global diversity and geography of soil fungi. Science, 346(6213), 1256688.

Thompson, L. R. et al. (2017). A communal catalogue reveals Earth’s multiscale microbial diversity. Nature.

Vinga, S. (2014). Information theory applications for biological sequence analysis. Briefings in Bioinformatics, 15(3), 376–389.

Xia, X. et al. (2003). An index of substitution saturation and its application. Molecular Phylogenetics and Evolution, 26(1), 1–7.

Yang, Z. (1994). Statistical properties of the maximum likelihood method of phylogenetic estimation and comparison with distance matrix methods. Syst. Biol., 43(3), 329–342.

Yilmaz, P. et al. (2014). The SILVA and "All-species Living Tree Project (LTP)" taxonomic frameworks. Nucleic Acids Research, 42(D1), D643–D648.

